# Arabidopsis EARLY FLOWERING 4 recruits EARLY FLOWERING 3 to the nucleus to facilitate gene repression

**DOI:** 10.64898/2025.12.12.688453

**Authors:** James Ronald, Seth J. Davis

## Abstract

The evening complex (EC) is a protein complex composed of ELF3, ELF4 and LUX that accumulates at dusk to repress gene expression. ELF3 functions as a scaffold around which the other EC components assemble, while LUX facilitates EC binding to DNA. Although ELF4 is essential for the activity of the EC, its precise role within the complex is unclear. Earlier studies indicated that mutations within the ELF4-binding domain of ELF3 reduced the nuclear accumulation of ELF3. However, it has remained unknown whether ELF4 directly contributes to these phenotypes. To investigate this further, we quantified the cellular and subnuclear distribution of Arabidopsis ELF3 across the evening in hypocotyl nuclei with or without a functional ELF4 allele. These results highlight a critical requirement for ELF4 in recruiting ELF3 to hypocotyl nuclei at dusk. Reduced nuclear accumulation in the *elf4* mutant correlated with a failure of ELF3 to repress gene expression and inhibit hypocotyl elongation, particularly under short-day conditions. We also observed that ELF4 was necessary to recruit ELF3 to foci in root nuclei, but these foci displayed different temporal properties from those in the hypocotyl. In summary, our results support a model in which ELF4 recruits ELF3 to the nucleus at dusk to enhance ELF3 repressive activity.

## Introduction

Circadian clocks are biological timekeeping mechanisms that facilitate the alignment of internal physiological and metabolic processes with the external environment by incorporating stimuli with predictable daily oscillations. In plants, the circadian clock controls daily processes such as growth and the response to abiotic and biotic stress, alongside key seasonal developmental milestones such as the timing of germination and the transition to flowering (Steed et al., 2021).

The plant circadian clock is composed of a series of interlocking transcriptional-translational feedback loops (TTFLs). At the center of these TTFLs are CIRCADIAN CLOCK ASSOCIATED 1 (CCA1), LATE ELONGATED HYPOCOTYL (LHY), and PSEUDO RESPONSE REGULATOR (PRR) TIMING OF CAB EXPRESSION 1 (TOC1) (McClung, 2019). CCA1/LHY and TOC1 mutually repress each other’s expression, restricting the phase of their expression to dawn and dusk respectively (Gendron et al., 2012, Kamioka et al., 2016). The expression profile of *CCA1/LHY* and *TOC1* are further regulated by additional day and evening loops. These include the day-phased *PRR9*, *PRR7*, and *PRR5* which are expressed sequentially from the early morning and repress *CCA1/LHY* expression (Wang et al., 2013). Finally, the evening complex (EC), composed of EARLY FLOWERING 3 (ELF3), ELF4, and LUX ARRHYTHMO, regulates *TOC1* and other circadian gene expression post dusk (Herrero et al., 2012, Ezer et al., 2017, Lee et al., 2019). Together, these regulatory connections shape the transcriptional architecture of the plant circadian clock.

So far, most studies have focused on resolving the transcriptional networks that form the plant circadian clock. However, over the last decade multiple studies have identified complex translational and post-translational mechanisms involved in timekeeping (Yan et al., 2021, Fan et al., 2022). In the animal and fungal circadian clock, controlling protein localisation is critical for circadian rhythms (Tataroğlu and Schafmeier, 2010). In plants, a few studies have highlighted that regulating protein localisation is also important for plant circadian rhythms (Wang et al., 2010, Kolmos et al., 2011, Kim et al., 2013a). However, the wider cellular dynamics of the plant clock has remained unclear.

ELF3 is a fundamental component of the plant circadian clock, with mutations in *elf3* resulting in circadian arrhythmia (Covington et al., 2001, Thines and Harmon, 2010). ELF3 is the scaffold around which LUX, ELF4 and chromatin remodelling enzymes assemble to form the EC (Nusinow et al., 2011, Herrero et al., 2012, Ezer et al., 2017, Park et al., 2019, Zha et al., 2020). ELF3 also functions independently of the EC, inhibiting transcription factors from binding to DNA and regulating protein stability (Yu et al., 2008, Nieto et al., 2015). ELF3 activity is thought to be exclusively nuclear and Arabidopsis ELF3 has a predicted NLS within the C-terminus (Liu et al., 2001). However, this NLS is only conserved amongst Brassicaceae. In monocots, ELF3 contain a distinct NLS in the middle region of the protein (Saito et al., 2012), while there is no conserved NLS amongst other ELF3 proteins. Therefore, other mechanisms must facilitate the nuclear localisation of ELF3.

Proteins lacking an NLS can localise to the nucleus through interactions with other proteins that contain an NLS, a process termed nuclear shuttling. Arabidopsis ELF4 is a small, single domain protein that intrinsically localises to the nucleus through a putative NLS in the N-terminus (Khanna et al., 2003). The ELF4-ELF3 interaction is conserved across multiple species (Lagercrantz et al., 2020, Cai et al., 2022, Zhao et al., 2022). For Arabidopsis ELF3, this interaction requires the box II domain termed ELF3-M (Herrero et al., 2012). Co-expressing Arabidopsis *ELF4* alongside Arabidopsis *ELF3*-*M* increased the nuclear localisation of ELF3M in tobacco mesophyll cells (Herrero et al., 2012). Furthermore, two independent point mutations in the ELF3-M region reduced the nuclear accumulation of ELF3 (Kolmos et al., 2011, Anwer et al., 2014). Although there were no overt physiological phenotypes caused by these mutations, both of these alleles independently shortened the circadian period (Kolmos et al., 2011, Anwer et al., 2014). ELF4 has also been demonstrated to promote the localisation of GIGANTEA (GI) to sub-nuclear bodies (Kim et al., 2013b). In contrast to ELF3, the recruitment of GI to these nuclear bodies was found to repress GI activity. Therefore, ELF4 seemingly regulates the cellular distribution of circadian proteins to control their functional activity.

So far, the role of ELF4 in facilitating the recruitment of ELF3 to the nucleus has only been shown in transient systems using a single domain of ELF3 or indirectly inferred through mutations within ELF3M (Anwer et al., 2014, Herrero et al., 2012, Kolmos et al., 2011). However, the ELF3M region mediates interactions with other proteins that could contribute to the nuclear localisation of ELF3 (Yu et al., 2008). To confirm that ELF4 directly regulates the nuclear accumulation of ELF3, we investigated the cellular and sub-nuclear distribution of ELF3 across an evening in the presence or absence of functional *ELF4* allele in Arabidopsis. These results demonstrate a critical requirement for ELF4 in facilitating ELF3 nuclear localisation but the importance of ELF4 was dependent on the tissue type. The recruitment of ELF3 to the nucleus by ELF4 was necessary to repress the expression of key EC circadian target genes. Furthermore, ELF4 was necessary for ELF3 to efficiently repress hypocotyl elongation, particularly under photoperiods with a shorter daylength. Together, these results confirm a role for ELF4 in promoting ELF3 nuclear and sub-nuclear localisation to facilitate ELF3 functional activity.

## Methods

### Plant lines and husbandry

All transgenics and mutant lines used in this work are in the Wassilewskija (Ws-2) background. The ELF3ox *elf3-4 (elf3-)* transgenic line, *elf3-4 LHY::LUC* and *elf3-4/elf4-1 LHY::LUC* mutants have all been described previously (Herrero et al., 2012). To generate the ELF3ox *elf3-4/elf4-1* mutant, the ELF3ox *elf3-4* line was crossed to the *elf3-4/elf4-1 LHY::LUC* mutant. A non-segregating ELF3ox *elf3-4/elf4-1 (elf3/elf4-) LHY::LUC* mutant was then identified in subsequent generations through genotyping.

### Growth conditions for microscopy

Seeds of either ELF3ox *elf3-* or ELF3ox *elf3/elf4-* were surface-sterilised and plated onto 1x MS media supplemented with 1.5% phytoagar, 0.25% sucrose and 0.5 g/L MES with a pH of 5.7. Seeds were stratified at 4°C for 3 days before being transferred to a short-day cabinet (8/16 photoperiod, constant temperature of 22°C) and grown vertically for six days. The imaging time course was then started at ZT7 on day 7. At each timepoint, a new plate was measured with 2 to 3 individual seedlings measured per timepoint per repeat. For timepoints after dusk, supplemental room lighting was restricted to a safe green light. All measurements per timepoint were concluded within an hour of the start time. The same microscope settings described in (Ronald et al., 2022) were used regardless of the context of the *elf4* allele. All data was made relative to the values of ELF3ox *elf3*- at ZT7. For each genotype and time point, the experiment was repeated on two separate occasions. The presented data is a combination of these two repeats.

### RT-qPCR

Seeds of the respective line were surface-sterilised and plated onto 1x MS media supplemented with 1.5% phytoagar, 0.25% sucrose and 0.5 g/L MES with a pH of 5.7. Seeds were stratified at 4°C for 3 days before being transferred to a short-day cabinet (8/16 photoperiod, constant temperature of 22°C). On day 7, ∼100 mg of seedlings from each respective genotype were harvested at ZT12 under a safe green light and subsequently snap-frozen in liquid nitrogen. RNA extractions, qPCR and subsequent data analysis was then performed as described before (Ronald et al., 2022).

### Hypocotyl Measurements

Seeds of the respective line were surface-sterilised and plated onto 1x MS media supplemented with 1.5% phytoagar, 0.25% sucrose and 0.5 g/L MES with a pH of 5.7. Seeds were stratified at 4°C for 3 – 4 days before being exposed to white light (25 µmol/m-2/s-1) for 6 hours to induce germination. Afterwards, the plates were transferred to a photoperiod chamber set to the appropriate daylength, a light intensity of 75 µmol/m-2/s-1 and a constant temperature of 22°C. Daylength started with 0 hours of light (constant darkness) and increased in four-hour intervals until 24 hours of light (constant light). Hypocotyl measurements were taken 7 days after stratification had ended, with plates harvested before dusk under each respective photoperiod. Two biological repeats were measured per photoperiod, with a minimum of 11 seedlings measured per repeat. The presented data is a combination of these repeats.

## Results

### ELF4 recruits ELF3 to the nucleus at dusk in hypocotyl cells

To investigate a possible role of ELF4 in directly regulating the cellular and sub-nuclear dynamics of ELF3, the *elf4-* mutant was introgressed into the previously described *35S::YFP:ELF3 elf3-* line (Herrero et al., 2012) to generate a homozygous *35S::YFP:ELF3 elf3/elf4-* double mutant. The cellular and sub-nuclear localisation of ELF3 in hypocotyl cells was then imaged in these two backgrounds. Seedlings of the respective construct were grown under a short-day (SD) photoperiod before imaging was started at ZT7 (1-hour before dusk) on day 6. Seedlings were then imaged at ZT8 and then every four hours following this until dawn on day 7 (ZT24).

ELF3 was localised between the cytoplasm and nucleus of hypocotyl nuclei at all timepoints and in both genetic backgrounds (**Figure 1A, 1C-D**). However, the extent of ELF3 nuclear localisation was dependent on both the time of day and the context of the *ELF4* allele. In a WT ELF4 background, the ELF3 nuclear signal increased between ZT7 and ZT8, before peaking at ZT12 (**Figure 1A, 1C**). There was then a sharp decline in the ELF3 nuclear signal at ZT16, before recovering across the remaining timepoints. These oscillations in ELF3 nuclear abundance were abrogated by the *elf4* mutation (**Figure 1A, 1D**). In the absence of *elf4*, the ELF3 nuclear signal displayed no change between ZT7 and ZT8 and failed to peak at ZT12. Instead, the nuclear levels of ELF3 declined across the early evening before partially recovering at ZT16. However, at no timepoint did ELF3 nuclear levels in the *elf4* background peak above those of WT and remained distributed between the nucleus and cytoplasm (**Figure 1A**). Together, these highlight an essential role for ELF4 in promoting ELF3 nuclear abundance, especially around the light to dark transition.

**Figure 1-.**
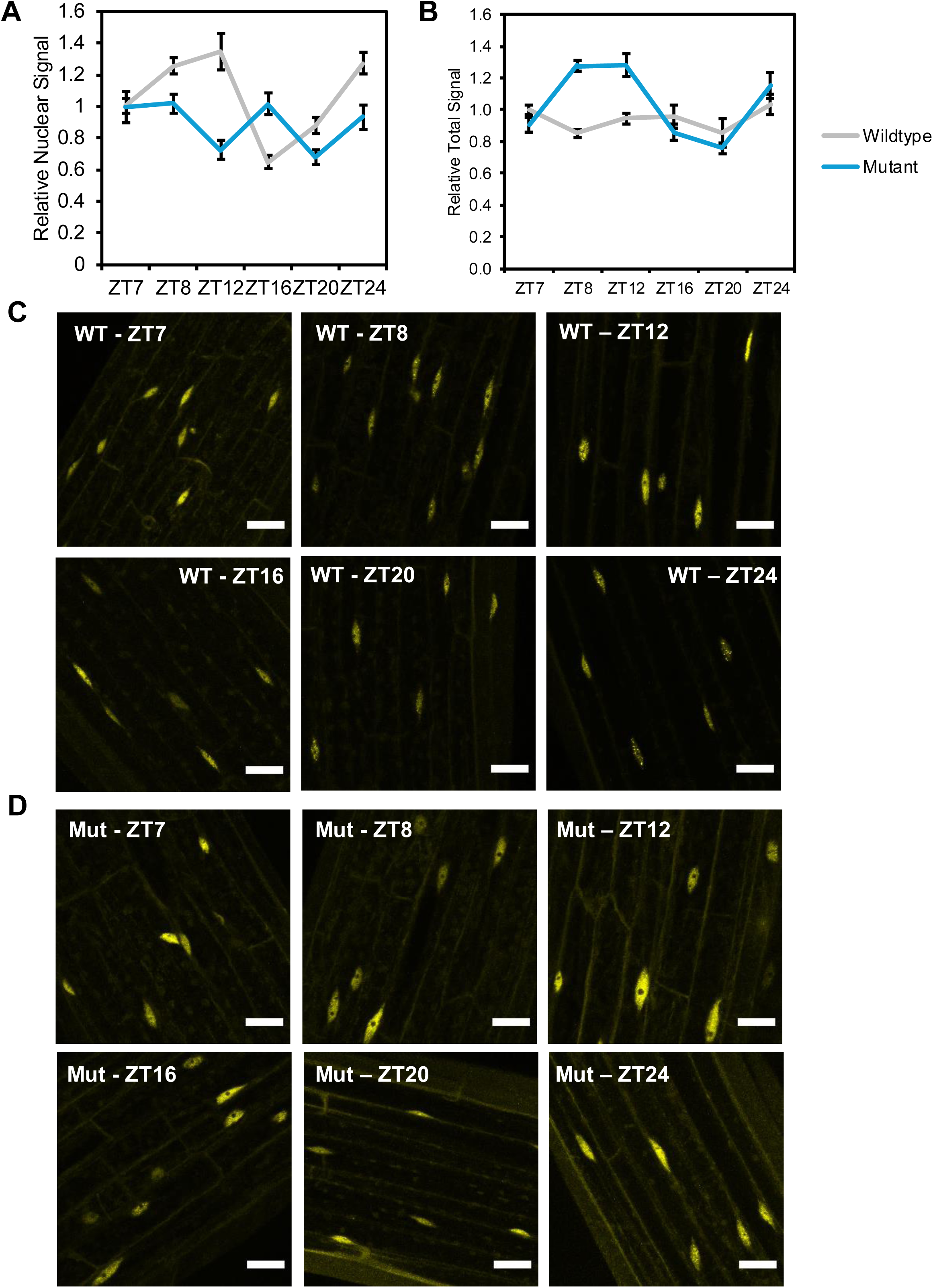
ELF3 nuclear localisation across time is dependent on ELF4. **(A)** The nuclear or **(B)** total signal of *35S::YFP:ELF3* across a short-day evening in either the *elf3-4* (wild type, grey) or *elf3-4/elf4-1* (mutant, blue) background. All data was normalised to the nuclear signal at ZT7 in the *elf3-4* background. Error bars are standard error of the mean. Representative images of **(C)** *elf3-4* or **(D)** *elf3-4/elf4-1* at each timepoint. Each timepoint was imaged on two separate occasions, with multiple independent seedlings measured each time. Scale bars are 25 μm.

To understand the relationship between nuclear abundance and protein levels, we used the YFP signal as an estimate for ELF3 protein abundance (**Figure 1B**). In a WT background, the YFP-ELF3 signal displayed an approximate 20% decline following lights off but then remain relatively stable across the night. In contrast, the YFP-ELF3 signal in the *elf4* background demonstrated larger changes across the timeseries (**Figure 1B**). For example, at ZT8 and ZT12, when the ELF3 nuclear levels were observed to decline, we found that YFP-ELF3 total signal increased in the *elf4* background. At the later timepoints, ELF3 signal in the *elf4-* background declined to levels comparable with WT. This suggests that nuclear localisation of ELF3 may be associated with reduced protein stability.

### ELF3 cellular dynamics are tissue-type dependent

ELF3 has been reported to form nuclear foci in multiple species at ambient and warm temperatures (Yu et al., 2008, Herrero et al., 2012, Anwer et al., 2014, Jung et al., 2020, Ronald et al., 2021, Ronald et al., 2022, Murcia et al., 2022). The ability of a protein to form sub-nuclear structures is tightly linked with protein concentration (Emenecker et al., 2021). As the nuclear levels of ELF3 were lower in the *elf4* background (**Figure 1B**), we hypothesised that the *elf4* mutation may also reduce ELF3 forming sub-nuclear foci.

We investigated ELF3 foci accumulation in hypocotyl nuclei across the same timeseries as in Figure 1. As reported before, ELF3 in the WT background formed nuclear foci at all timepoints (**Figure 2**). However, the number and morphology of the foci was dependent on the time of day. Unlike with the nuclear accumulation of ELF3, there was no change in the number or morphology of foci between ZT7 and ZT8 in the WT background. Maximal foci accumulation occurred at ZT12 (**Figure 2A-B**), correlating with the maximal nuclear accumulation of ELF3, while the fewest foci were observed at ZT16 when the nuclear accumulation of ELF3 was at its lowest. Foci were largest and most distinguishable from the nucleoplasmic signal at ZT12 and ZT24 (**Figure 2B**). In the *elf4* background, these time-of-day effects on foci formation were completely lost (**Figure 2A, 2C**). At all timepoints, there were fewer foci in the *elf4* nuclei compared to WT. Furthermore, when ELF3 foci in the *elf4* background were observed, these were smaller and less bright than those observed in WT nuclei (**Figure 2C**). Together, these results illustrate a requirement for ELF4 to mediate ELF3 localisation into nuclear foci.

**Figure 2-.**
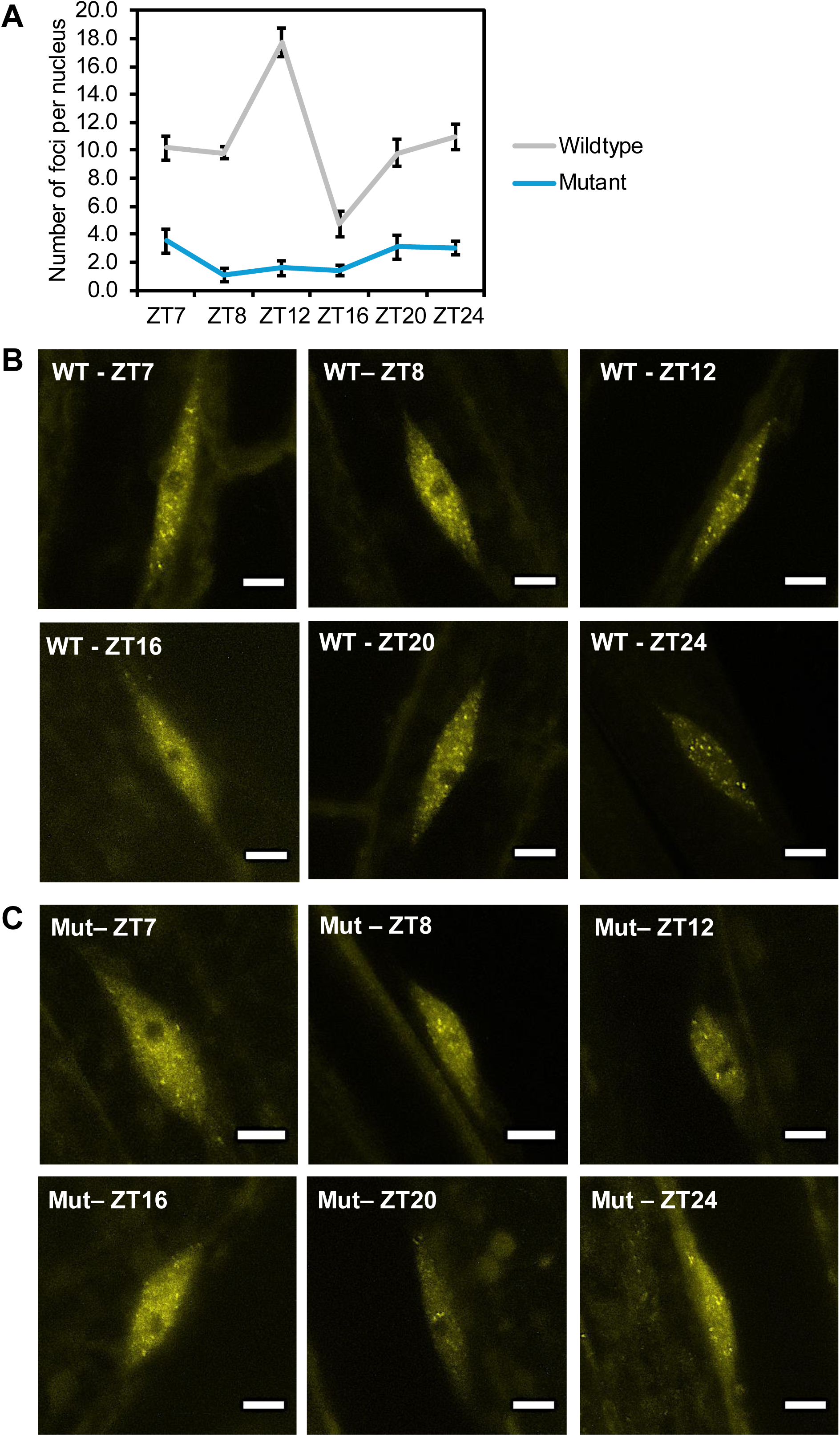
ELF4 facilitates the sub-nuclear localisation of ELF3. **(A)** The number of ELF3 foci per nuclei across a short-day evening in either the *35S::YFP:ELF3 elf3-4* (wild type, grey) or *35S::YFP:ELF3 elf3-4/elf4-1* (mutant, blue) background. Representative images of ELF3 foci in the **(C)** *elf3-4* or **(D)** *elf3-4/elf4-1* background at each timepoint. Each timepoint was imaged twice, with multiple independent seedlings measured each time. All imaging was performed on two separate occasions. Scale bars are 5 μm.

### ELF3 localisation to foci highlights tissue-specific dynamics

We next investigated whether ELF3 sub-nuclear dynamics were similar in root nuclei. Here we focused on three timepoints, ZT7, ZT8 and ZT12. As we have reported previously (Ronald et al., 2021), foci in root nuclei were morphologically different than their counterparts in hypocotyl tissue, appearing larger and more distinguishable from the nucleoplasmic signal (**Figure 3A**). The number of foci in root nuclei did not show the same time of day effects as observed in hypocotyl nuclei (**Figure 2A, 3B**) but the number of foci were more variable in root nuclei (**Figure 3B**). Sorting nuclei by the time the image was taken revealed a strong increase in the number of foci leading up to and across the light/dark boundary in root nuclei (**Supplementary Figure 1A, 1C**). This behaviour was not observed in hypocotyl nuclei (**Supplementary Figure 1B, 1D**). Thus, ELF3 foci in root nuclei have different temporal properties than in hypocotyls, possibly reflecting tissue-specific dynamics of the plant circadian clock.

**Figure 3-.**
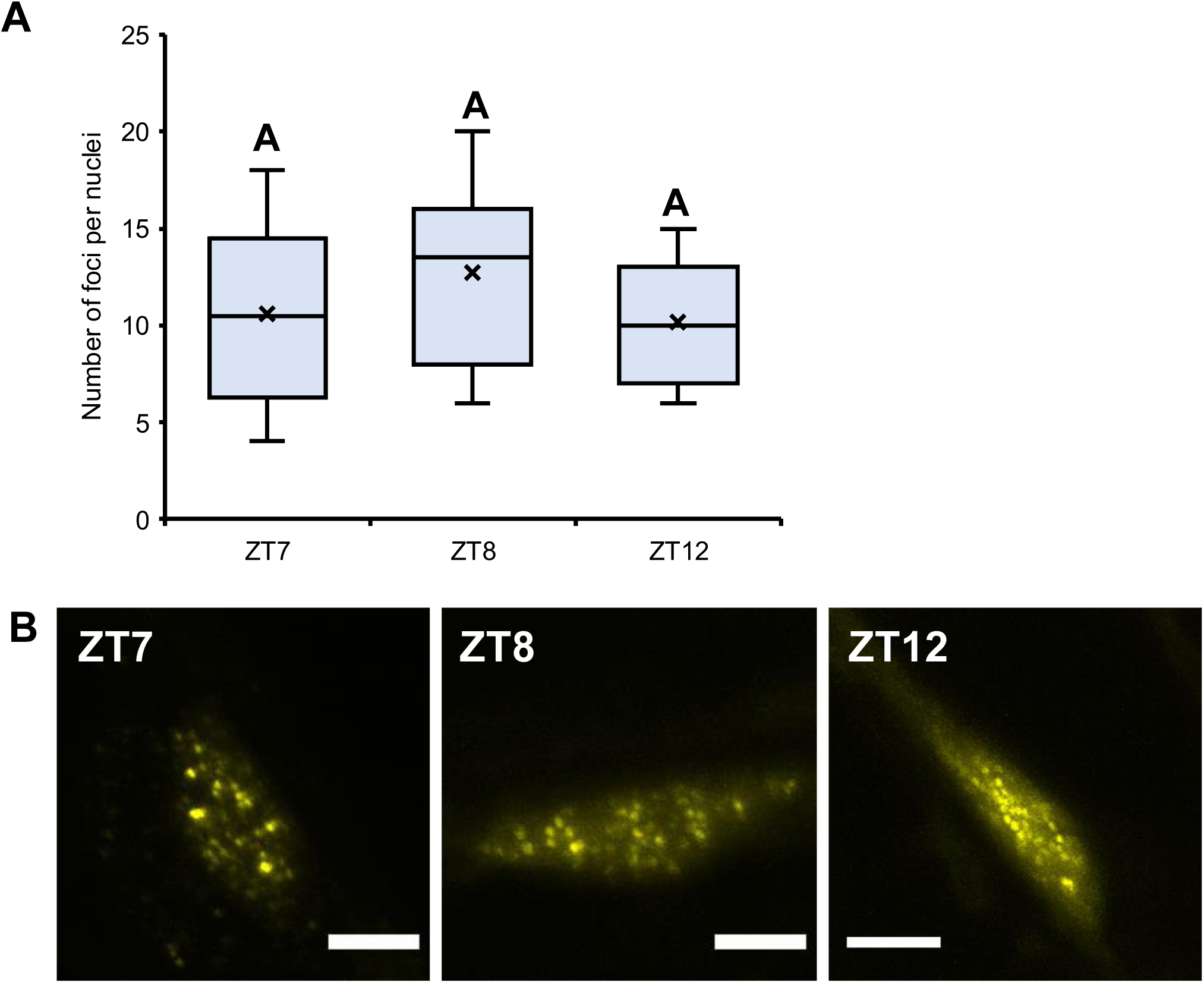
ELF3 foci display different temporal properties in root nuclei. **(A)** The localisation of ELF3 to foci in root nuclei at ZT7, ZT8 and ZT12 (short-day photoperiod). **(B)** Representative images of ELF3 foci at the respective timepoints. Each timepoint was imaged on two separate occasions, with multiple independent seedlings measured each time. Letters indicate no statistical significance between the respective timepoints as determined by an ANOVA with a tukey-HSD post hoc test.

ELF4 is a mobile protein that moves between the hypocotyl and root in a temperature-dependent manner (Chen et al., 2020). Thus, we investigated whether ELF4 is also regulating ELF3 sub-nuclear dynamics in root tissue. As in hypocotyl tissue, mutating *elf4* impaired the ability of ELF3 to form nuclear condensates **(Supplementary Figure 2A)**, consistent with our previous results (Ronald et al., 2021). However, the impact of the *elf4* mutation was weaker in root nuclei compared to hypocotyl nuclei (**Supplementary Figure 2B**). There was also less variation in the number of foci per root nuclei at both ZT7 and ZT8 in the *elf4* mutant background and subsequently there was no clear pattern of foci accumulating around the light/dark boundary as observed in WT nuclei (**Supplementary Figure 2C-D**). These results indicate that ELF4 may facilitate the recruitment of ELF3 to foci across the light-dark boundary in root nuclei.

### ELF4 is required for ELF3 to repress gene expression

To understand how changes in the cellular localisation of ELF3 influenced ELF3 activity, we firstly compared the expression of ELF3/EC targets in the *ELF3ox* background with or without a functional *ELF4* allele at ZT12. ZT12 was chosen as this timepoint had the largest difference in the nuclear and sub-nuclear accumulation of ELF3 between the two respective backgrounds (**Figure 1, 2**). There was no change in the expression *LUX*, *TOC1* or *GI* expression in the *ELF3ox elf3-* background compared to wild type (WT) plants at ZT12 (**Figure 4**). These results support earlier results that found no strong repressive effect of *ELF3ox* at ZT12 (Nieto et al., 2015). In contrast, ELF3ox in the *elf3/elf4-*background failed to repress the expression of *GI* or *LUX* compared to WT or *ELF3ox elf3-*(**Figure 4A-B**). The expression of *TOC1* was also elevated in the ELF3ox *elf4-* background, although this effect was weaker than for *GI* or *LUX* (**Figure 4C**). Combined, these results suggest that ELF4 is required for ELF3 to repress the expression of *GI, LUX* and *TOC1* in the evening.

**Figure 4-.**
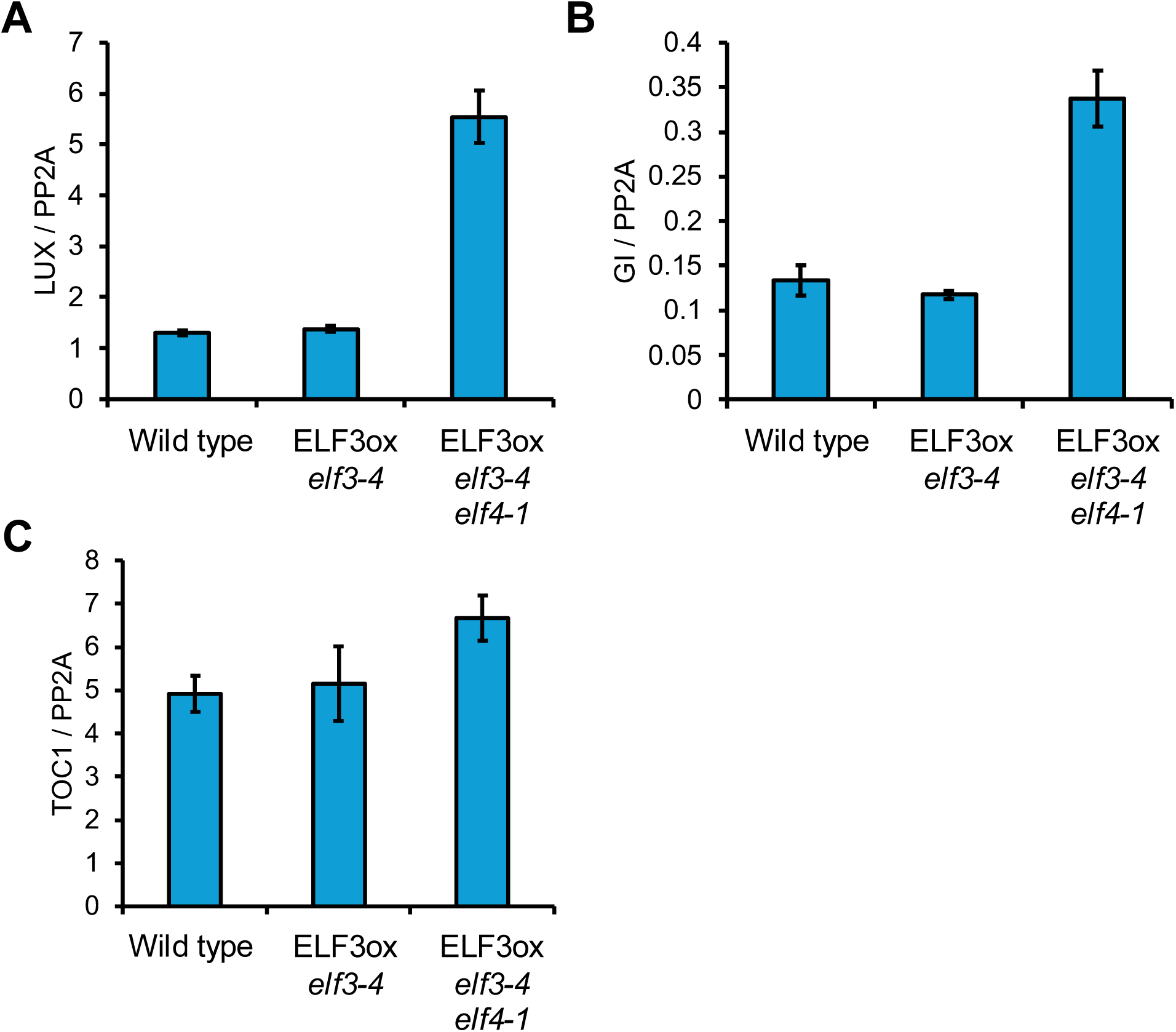
ELF4 is required for the repression of Evening Complex Targets. The expression of (A) *LUX ARRYTHMO* (*LUX*), (B) *GIGANTEA* (*GI*) or (C) *TIMING OF CAB EXPRESSION 1* (*TOC1*) in Ws-2 *LHY::LUC* (WT), *ELF3ox elf3-4* or ELF3ox *elf3-4/elf4-1* background at ZT12 (short-day photoperiod). Expression of the respective gene is normalised to *PROTEIN PHOSPHATASE 2A*. The presented data is an average of three technical repeats, with error bars representing standard deviation. All experiments were repeated with two separate samples with similar results observed on both occasions.

### ELF4 facilitates ELF3 repression of hypocotyl elongation under shorter-day photoperiods

To investigate the importance of ELF4 in facilitating ELF3’s regulation of plant development, we measured hypocotyl elongation in wild type (WT), *elf3-*, *elf3-/elf4-*, 35S*::YFP:ELF3* (*elf3-*) and *35S::YFP:ELF3* (*elf3/elf4-*) seedlings grown across a range of photoperiods. As described before (Anwer et al., 2020), the hypocotyl length of WT seedlings was inversely related to the length of the photoperiod. As the duration of the photoperiod increased, hypocotyl elongation in WT seedlings was progressively inhibited (**Figure 5**). The *elf3-* single and *elf3/elf4-* double mutant had longer hypocotyls than WT seedlings under all diel photoperiods as reported previously but this particularly enhanced under short- and neutral day photoperiods (**Figure 5**). There was no pronounced enhancement of the *elf3-/elf4-* phenotype relative to *elf3-*.

**Figure 5-.**
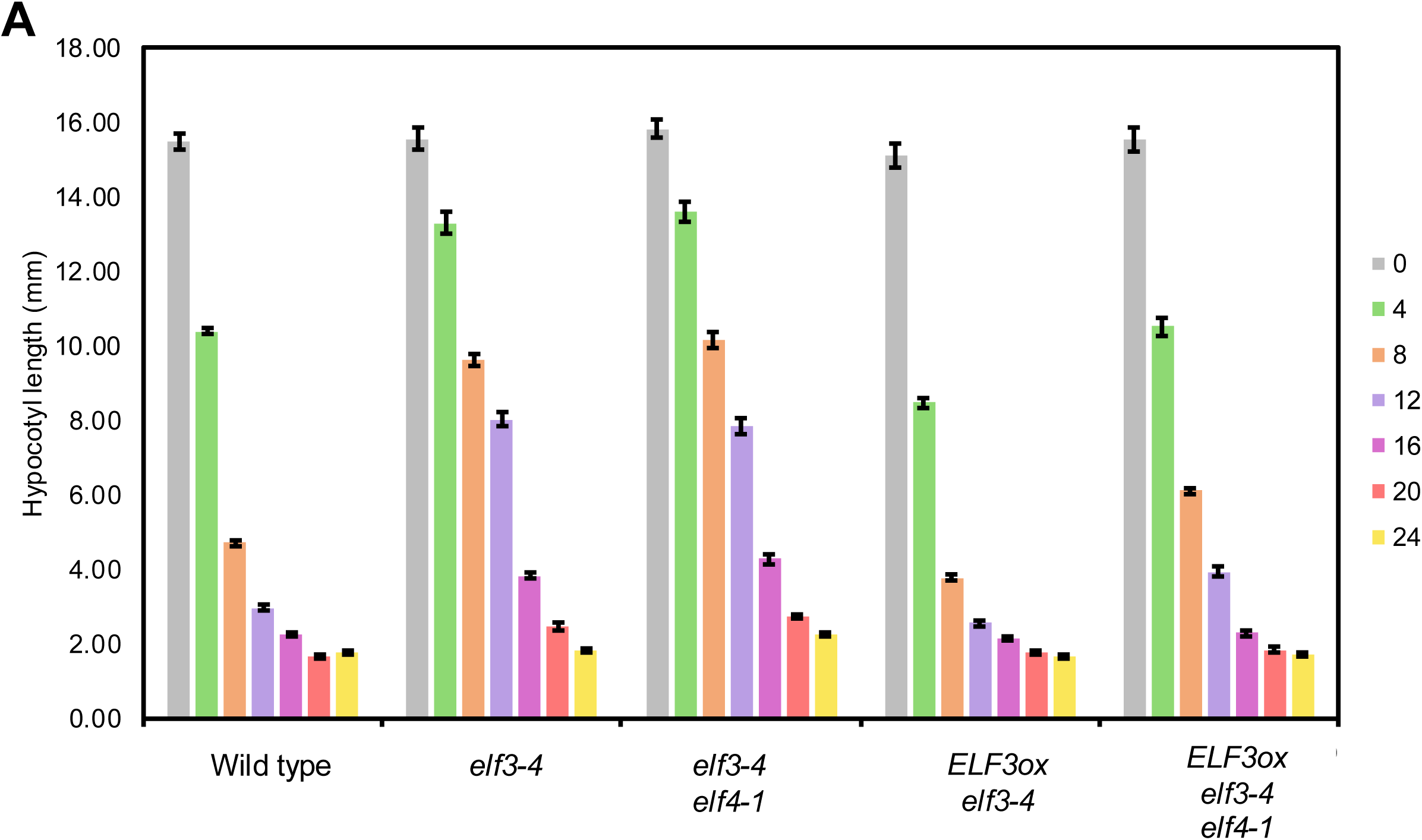
ELF4 facilitates ELF3 repression of hypocotyl development under shorter day lengths. Hypocotyl length was measured in Ws-2 *LHY::LUC*, *elf3-4 LHY::LUC*, *elf3-4/elf4-1 LHY::LUC*, *ELF3ox elf3-4 LHY::LUC* and *ELF3ox elf3-4/elf4-1 LHY::LUC* across a range of photoperiods from no light (0) to constant light (24). Temperature was 22°C for all photoperiods. Each genotype per photoperiod had a minimum n of 10 seedlings. All experiments were repeated twice, with no statistical significance between each repeat. The presented data is a combination of both experimental repeats. Error bars represent standard error of the mean.

The *elf3-* hypocotyl phenotype was fully rescued by introducing *ELF3ox* under all photoperiods (**Figure 5**). Furthermore, in photoperiods with a light phase that was less than 16-hours in length, *ELF3ox* hypocotyls were shorter than WT seedlings. In contrast, *ELF3ox* could only partially rescue the *elf3/elf4-* phenotype. Under short-days (8/16) and neutral days (12/12), the hypocotyl of *ELF3ox* was longer than WT. The hypocotyl length of *ELF3ox* under these photoperiods was not as long as the *elf3/elf4-* double mutant, indicating ELF3 retained some activity in the absence of *elf4* (**Figure 5**). For photoperiods with 16 or more hours of light, *ELF3ox elf3/elf4-* had a similar hypocotyl length to WT and *ELF3ox* e*lf3-*. This indicates that ELF4 may have a critical role in facilitating ELF3 inhibition of hypocotyl elongation under photoperiods with a longer dark phase.

## Discussion

Precisely controlling where and when ELF3 is localising in the cell is an emerging mechanism that is seemingly key for controlling ELF3 activity (Kolmos et al., 2011, Anwer et al., 2014, Jung et al., 2020, Ronald et al., 2021, Murcia et al., 2022). Here, we have found that ELF3 nuclear and sub-nuclear distribution in hypocotyl cells displays clear time of day effects (**Figure 1, 2**), although these are weaker for foci accumulation. Both nuclear localisation and sub-nuclear foci accumulation peaked in the early evening, approximately four hours after dusk, before declining in the late evening and then recovering towards dawn. Mutating *elf4* strongly impaired the nuclear localisation of ELF3 and subsequent ability to undergo phase separation. As ELF4 has cytoplasmic localisation in Arabidopsis (Herrero et al., 2012), it is possibly that ELF4 could directly shuttle ELF3 to the nucleus The structure and function of ELF4 is seemingly conserved across evolution, with Chlamydomonas ELF4 sharing the same domain structure as Arabidopsis ELF4 and can rescue the Arabidopsis *elf4* flowering phenotype (Zhao et al., 2019). Thus, ELF4 recruitment of ELF3 to the nucleus may represent the primary mechanism for how ELF3 localises to the nucleus, with the evolution of the NLS a unique feature in certain plant species.

Why ELF3 in Arabidopsis may rely on two mechanisms of nuclear importation exist is unclear. The requirement of ELF4 to facilitate the nuclear localisation of ELF3 was most apparent in the early evening (**Figure 1**). This correlates with the timing for when the EC starts to assemble (Nusinow et al., 2011) and when plants are undergoing large-scale transcriptional changes in response to the light/dark transition (Ezer et al., 2017). Thus, the recruitment of ELF3 to the nucleus by ELF4 at dusk could enable rapid assemblage of the EC. This in turn would facilitate the repression of genes associated with photosynthetic or other light-sensitive processes to switch the plants into ‘night-mode’. In contrast, the NLS-dependent pathway could maintain a steady-state level of nuclear ELF3 across the evening to enable non-EC dependent functions that have been described elsewhere (Nieto et al., 2015, Yu et al., 2008). The recruitment of ELF3 to the nucleus by ELF4 physiologically seems to be most critical under a shorter photoperiod (**Figure 5**). It will be important to investigate this further.

ELF4 was also necessary to facilitate the recruitment of ELF3 to nuclear foci in hypocotyl and root nuclei (**Figure 2, Supplementary Figure 2**). Whether this is a direct role of ELF4 in mediating condensation of ELF3 or is an indirect effect caused by increasing ELF3 nuclear levels is uncertain. The ability of proteins to undergo condensation is tightly linked with their protein abundance (Emenecker et al., 2021), so it is not possible to untangle these two responses currently. Although the *elf4* mutation did impair foci accumulation in root nuclei, this phenotype was weaker root than hypocotyl nuclei. This is consistent with emerging reports on the EC and the wider circadian clock system being dependent on the tissue type (Davis et al., 2022). Combined, our results highlight a critical role for ELF4 in regulating the cellular dynamics of ELF3.

The role of foci in ELF3 activity continues to remain unclear. Previous studies have suggested that ELF3 forms foci at ambient temperatures for an unclear functional purpose (Herrero et al., 2012, Anwer et al., 2014, Ronald et al., 2022, Murcia et al., 2022). Separately, foci have been proposed to be sites of inactive ELF3 formed upon exposure to elevated temperature (Jung et al., 2020). Here, we found that ELF3 foci formation peaked at ZT12 (**Figure 2**). However, there was no change in the expression of *TOC1*, *LUX* or *GI* in the *ELF3ox* background relative to WT at ZT12 **(Figure 4)**. The absence of ELF3 repressing gene expression at ZT12 is consistent with earlier reports (Nieto et al., 2015) and ChIP-seq data that indicates LUX begins to dissemble from chromatin at ZT12 (Ezer et al., 2017). Together, this would suggest that the association of ELF3 to foci may not be indicative of sites of ELF3/EC activity.

If condensates are not sites of gene repression, then it continues to be unclear what roles ELF3 foci are fulfilling at ambient temperatures. One possible function of ELF3 foci could be storage sites to protect against proteolytic degradation by CONSTITUTIVE PHOTOMORPHOGENIC1 (COP1) (Yu et al., 2008). Supporting this, the nuclear levels of ELF3 in the *elf4* background were elevated during the early evening when ELF3 in this background was more localised to the cytosol (**Figure 1**). Foci functioning as storage sites has been previously proposed (Murcia et al., 2022). Finally, we have found that root foci appear morphologically different from hypocotyl foci and display different temporal properties (**Figure 2, 3**). Thus, it is possible that ELF3 forms multiple species of nuclear bodies, dependent on both environmental factors (light, temperature and time) and intrinsic factors such as protein concentration and protein-protein interactions.

## Acknowledgements

We thank the Horticulture team within the Biology Department for their support with plant care and facilities maintenance. We also thank the Imaging and Cytometry, and Genomics teams within the York Technology Facility for their assistance with the experiments. JR was funded by a BBSRC studentship 1792522 and a BBSRC fellowship BB/Z514998/1. SJD was funded by a BBSRC responsive mode grant BB/N018540/1.

---

**Supplementary Figure 1-.**
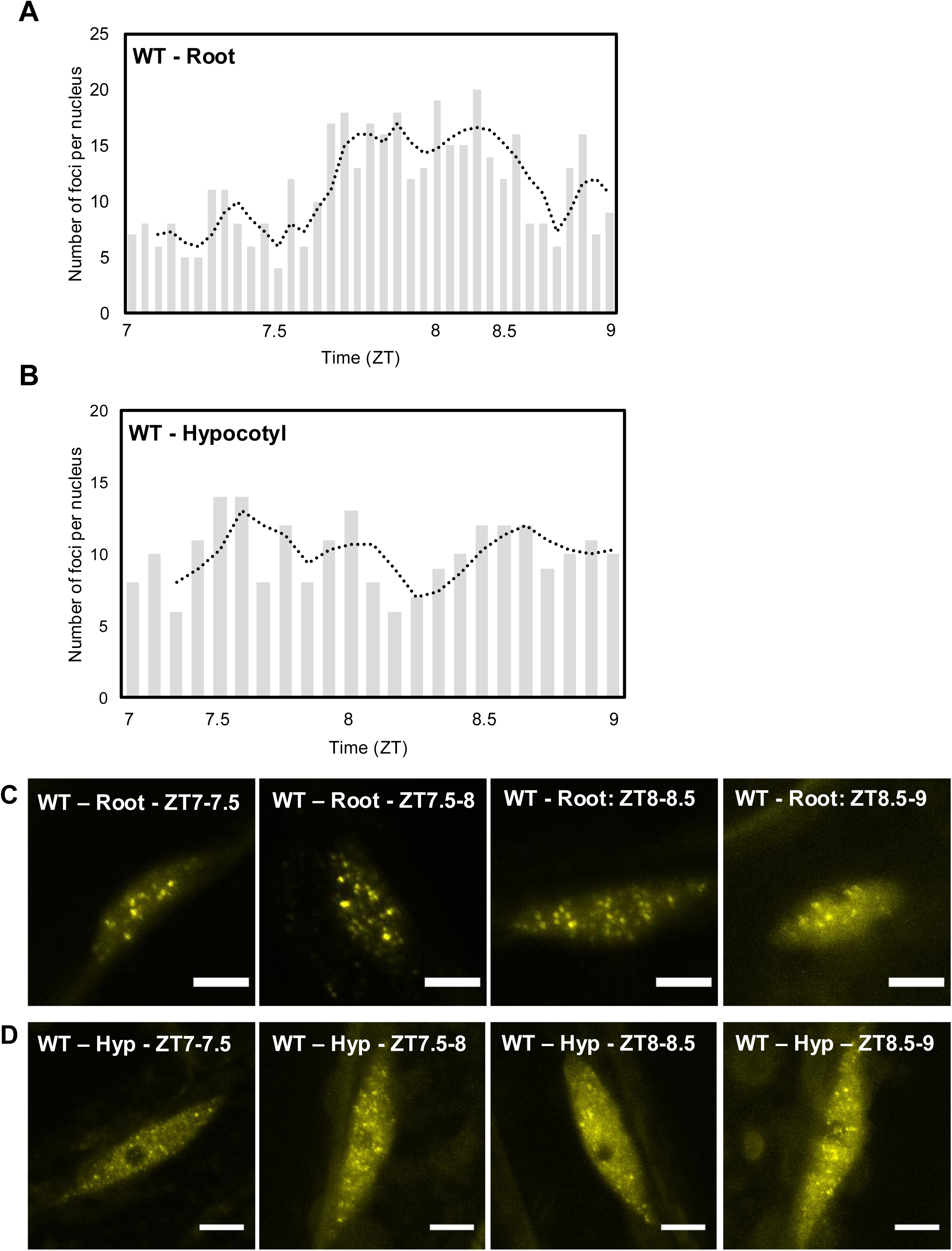
ELF3 foci in root nuclei accumulate at the light/dark boundary. **(A-C)** The formation of ELF3 foci in (A) root or (B) hypocotyl nuclei across ZT7 and ZT8 (short-day photoperiod) in the *ELF3ox* (*elf3-4*, termed WT) background. Each bar indicates an individual nucleus sorted by the time (ZT) each image was taken, ZT8 is dusk. Dashed line is a rolling average of 3. Data is a collation of two separate experiments. **(C, D)** Representative images of ELF3ox (*elf3-4*, termed WT) in **(C)** roots and **(D)** hypocotyls (hyp). Scale bars are 5 μm.

**Supplementary Figure 2-.**
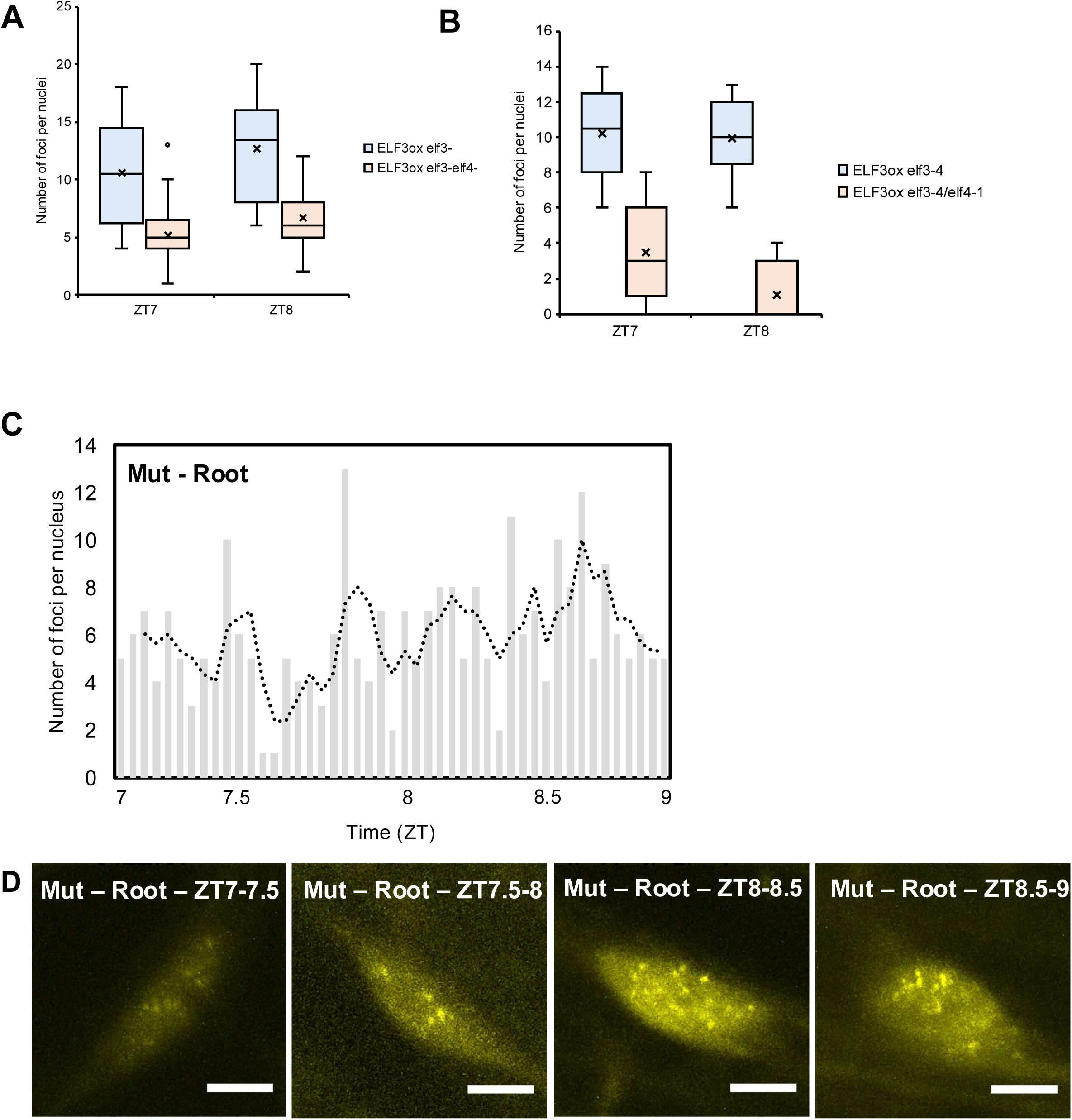
ELF4 has a weaker effect on foci formation in root nuclei compared to hypocotyl nuclei. **(A, B)** The number of ELF3 foci in the ELF3ox *elf3-4* or ELF3ox *elf3-4/elf4-1* background in (A) root nuclei or (B) hypocotyl nuclei at ZT7, ZT8 and ZT12. The hypocotyl data is the same as figure 2. The ELF3ox *elf3-4* root data is the same as Figure 3. All imaging was repeated twice on two separate occasions with multiple samples each time. The presented data is a combination of those two experiments. **(C)** The number of ELF3 foci in the ELF3ox (*elf3-4/elf4-1*) background across ZT7 – ZT9 in root nuclei. Each bar indicates an individual nucleus sorted by the time (ZT) each image was taken, ZT8 is dusk. Dashed line is a rolling average of 3. **(D)** Representative images of (B). Scale bars are 5 μm.

